# Quantifying the deformability of malaria-infected red blood cells using deep learning trained on synthetic cells

**DOI:** 10.1101/2022.12.16.520724

**Authors:** Daniel T. Rademaker, Joshua J. Koopmans, Gwendolyn M.S.M. Thyen, Aigars Piruska, Wilhelm T.S. Huck, Gert Vriend, Peter A.C. ‘t Hoen, Taco W.A. Kooij, Martijn A. Huynen, Nicholas I. Proellochs

**Affiliations:** Center for Molecular and Biomolecular Informatics, Radboud University Medical Center, 6525 GA Nijmegen, The Netherlands; Radboud Center for Infectious Diseases, Medical Microbiology, Radboud University Medical Center, 6525 GA Nijmegen, The Netherlands; Institute for Molecules and Materials, Radboud University, 6525 AJ Nijmegen, The Netherlands; Baco Institute for Protein Science, Mindoro 5201, Philippines

**Keywords:** Cell tracking, Deep Learning, malaria, Plasmodium falciparum, deformity index

## Abstract

Several haematologic diseases, including malaria, diabetes, and sickle cell anaemia, result in a reduced red blood cell deformability. This deformability can be measured using a microfluidic device with channels of varying width. Nevertheless, it is challenging to algorithmically recognise large numbers of red blood cells and quantify their deformability from image data. Deep learning has become the method of choice to handle noisy and complex image data. However, it requires a significant amount of labelled data to train the neural networks. By creating images of cells and mimicking noise and plasticity in those images, we generate synthetic data to train a network to detect and segment red blood cells from video-recordings, without the need for manually annotated labels. Using this new method, we uncover significant differences between the deformability of RBCs infected with different strains of *Plasmodium falciparum*, providing clues to the variation in virulence of these strains.

## 1. Introduction

The malaria parasite, which in 2020 caused an estimated 627,000 deaths worldwide[1], spends part of its life cycle in human red blood cells (RBCs). In the most deadly of malaria parasites, Plasmodium falciparum, the intraerythrocytic developmental cycle lasts 48 hours, comprising the ring-, trophozoite-, and schizont-stages. During these stages, the parasite remodels the RBC by setting up an extensive export process, which involves approximately 10% of the parasite’s proteome[2]–[4]. As a result, the infected RBC (iRBC) becomes sticky and hyper-rigid. While these RBC modifications are critical for parasite survival, the stickiness and rigidity are also a cornerstone of virulence and, as such, they have been the topic of great interest in malaria research[2]–[4]. To study variations in rigidity, e.g., during the intraerythrocytic developmental cycle, or between different P. falciparum strains, hundreds to thousands of cells must be analysed to create statistically meaningful data. Progress in studying these RBC morphological changes, particularly in membrane mechanics, has been hampered by the critical lack of a suitable assay to measure the changes in rigidity in a high throughput and accurate manner[5], [6]. However, recent advances in microfluidic devices allow measuring RBC mechanics in large quantities[7].

Microfluidic devices (MDs) are tools to manipulate fluid on a scale from a few microns to a few hundred microns. Feature size in MDs is commensurate with individual cell size, thus making MDs useful for cell analysis[8]. MDs provide a low-cost, high-throughput method to study RBCs under varying conditions[7], [9]. For example, the effect of small blood vessels on RBCs can be emulated with the use of narrow constriction channels. Comparing the deformity index (DI) (i.e., the shape deformity) of an RBC before and after the tight passage then yields a deformity index difference (ΔDI) value per RBC indicating cell deformability. High-speed microscope imaging records the events in such MDs, producing movies that can easily contain thousands of cells. Manually evaluating such a large number of cells, though, is a daunting task. One study investigating the effects of blood bank storage on RBC deformability partly automated some of the MD-video processing using conventional image processing techniques[10]. However, in general, these techniques are limited to preprogramed solutions[11] and cannot handle unexpected events like motion blur, translucent cell edges, colliding cells, air bubbles, unusual cell shapes, background noise, MD artefacts, floating cell debris, cell-clusters, and illumination variations (Figure 1 depicts an example of a typical video frame). As a result, MD-video processing still requires time-consuming human labour to predefine all the exceptions and design algorithms to handle them. An example of this, is the work of Saadat et al.[12], who despite reporting tremendous success in determining the mechanical properties of RBCs in high-throughput, confirmed that image noise was detrimental for cell tracking and shape determination and were unable to handle abnormally shaped cells. Since all these factors are prevalent in iRBCs MD videos, a different approach should be considered. Machine learning, and in particular deep learning (DL) may be a more effective strategy to process the image data, including handling the many challenging technical and biological variations.

**Figure 1.**
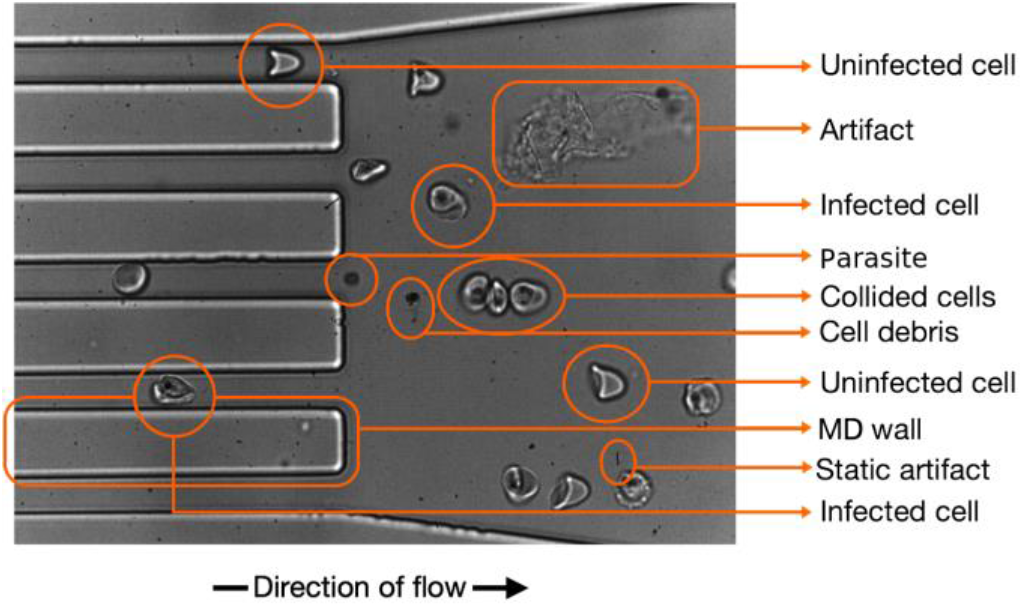
Topography of a typical microfluidic video frame. A typical video frame from a microfluidic device contains many objects other than the RBCs of interest. These non-cell objects, such as cell debris, microfluidic device walls, or other artefacts, in combination with varying focus and lighting conditions, interfere with the segmentation and tracking processes.

The success of many recent DL projects can be attributed to the combination of clever neural architecture design and automated feature extraction from large datasets[13]. In particular, the Convolutional Neural Network (CNN) architecture performs well on image data and has been shown to solve many complex image data problems with high accuracy[14]–[17]. Most relevant here is that it has shown great results in image segmentation —the per pixel labelling of images— under noisy conditions[11], [18], which in principle makes it an ideal candidate for handling MD video data. Given that DL is a data-driven approach, its performance directly depends on the quality and quantity of the data trained on. However, many scientific disciplines lack the amount of labelled data required to power DL algorithms. The use of synthetic data, which is data created by computer algorithms instead of collected from the real world, is one potential solution that is gaining traction in the DL community[19]. It provides a source of data that is similar to real-world data but with the benefits of having consistent labels and complete control over all characteristics of the data.

In this study, we created a robust method to automatically track and calculate DIs of RBCs from noisy MD videos with the goal of investigating the differences in RBC deformability between iRBCs of the geographically distinct Plasmodium falciparum strains K1[20] and NF135[21, p. 135]. We designed a CNN architecture with two output layers to translate the noisy videoframes into cell shapes and cell locations. We avoided the need for annotating a diverse, carefully labelled dataset of RBCs by training the CNN on synthetic data instead. We also created a Python programme that accepts the CNN outputs to extract the individual RBC journeys through the MD, remove colliding cells, and calculate cell ΔDIs. The extracted cell journeys were classified by a domain expert by stage of RBC infection. We confirm that there is a highly significant difference in RBC deformability between uninfected and early-stage infected RBCs, as well as a general trend of decreased deformability between the consecutive infection stages. Unexpectedly, we discovered a significant difference in deformability between the K1 and the NF135 strains at all stages of infection.

## 2. Results

### Generating synthetic data RBC images

We generated a synthetic training dataset of 10,000 image patches of 100×100 pixels that each could include RBCs, MD walls, and/or various artefacts. We here describe conceptually how this was done; see the Methods section for a more detailed description. RBCs in our experimental setting adopted either a round (Figure 2a) or a bullet-like shape (Figure 2c). Although the real-world data of RBCs and their background appear intricate (Figures 2a, 2c, and 2e), we were able to approximate them reasonably well with just a limited set of simple rules (Figures 2b, 2d, and 2f). For each of the synthetic columns b, d, and f, we used the same procedure to generate the diversity shown, i.e., all generated images started from circles, lines, or both. We warped these basic forms in a variety of ways using elastic-deformation, a popular technique for augmenting images[22]. To generate natural-looking textures, we used the Simplex-noise function, a function that produces gradient noise patterns that are commonly used in the game and film industries[23]. Doing so, we were able to generate the background (Figure 2f) and inner cell textures (Figure 2b). The noise patterns were also utilised as interpolation ratio matrices, which enabled us to interpolate between simple textures to create more complex ones, to make cells partially translucent, and to add a colour gradient to the background. In addition to the cell textures, we added dark spot(s) to a fraction of the cells to simulate the presence of parasites (Figure 2b). Finally, we used a blur technique to reduce harsh edges, thereby making the images appear more natural. The backgrounds were created in duplicate. One containing all objects, and one without any transient objects but with altered colour intensities to make the neural network robust for differences in lightning.

**Figure 2.**
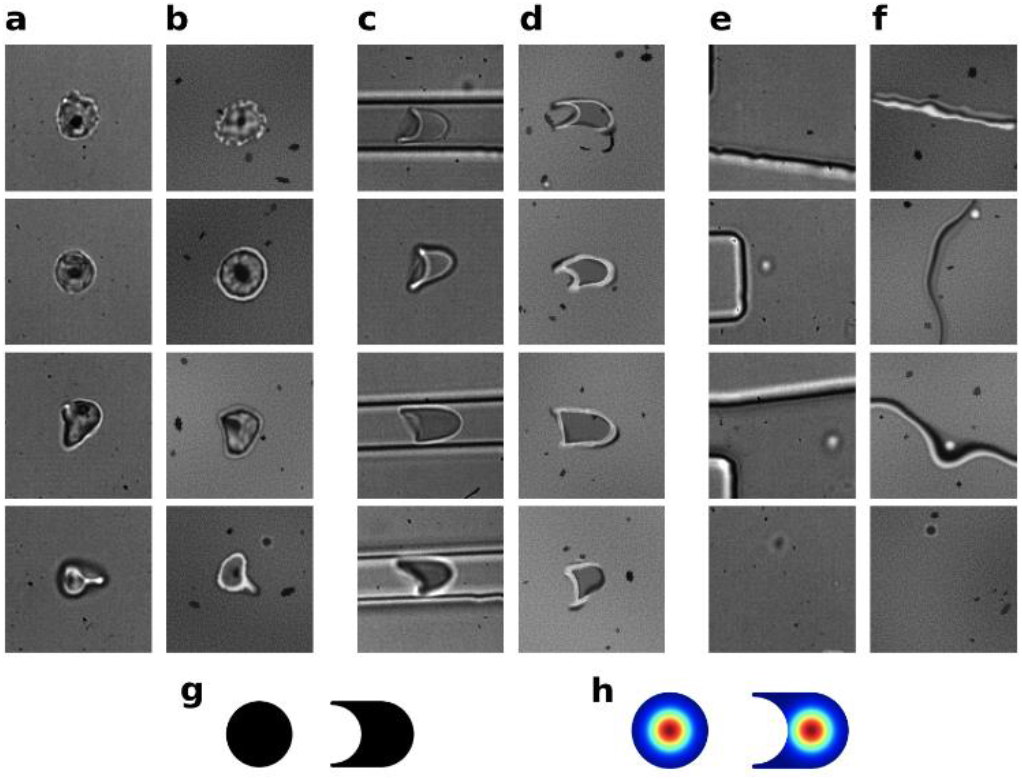
Synthetic and real-world example images. (a) Variation of real-world ‘round’ shaped RBCs; (c) same as (a), but with ‘bullet’ shaped RBCs; (e) same as (a) and (c) but with focus on background variations. (b), (d), and (f) are manually selected images from the synthetic data that are similar to the corresponding images in (a), (c), and (e), respectively. Labels are given in two ways: 1) the segmentation, indicating the probability per pixel that it belongs to an RBC, as illustrated in (g); and 2) a Gaussian distribution indicating the probability of pixels belonging to the centre of a RBC, as illustrated in (h).

All images were of course perfectly annotated because of the synthetic nature of the data. Each image had labels describing the cell’s form (Figure 2g) and labels describing the cell’s position (Figure 2h). The form labels contained pixel-per-pixel binary segmentation values indicating the probability that a pixel belonged to an RBC. For the cell location labels, each pixel was assigned the probability of being an RBC centre, modelled as a 2D Gaussian distribution with its peak at the cell centres (illustrated as a heat-map in Figure 2h). The synthetic images were significantly smaller than the video frames, as can be appreciated by comparing Figure 1 with Figure 2. This allowed for a much smaller and more efficient CNN to be trained on while still containing enough context for the CNN to perform well. An interesting finding during testing was that exaggerating the objects’ features helped the neural network generalise better. Making objects brighter or darker than they appeared in the videos, or having objects with more deformity than seen in the real data, for example, aided in making the neural network more robust.

### Red blood cell tracking and segmentation results

Training the CNN on the synthetic data allowed us to recognise ‘real’ RBCs from the MD image data. The CNN produced two kinds of output: one showing where the cells were and the other showing what shape they had. Using the CNN’s location outputs, a Python script tracked the cells throughout the video. Figure 3 depicts some of the resulting cell journeys, which were created by combining crops from each frame in which the cell was present. The accurate segmentation, which entails marking each pixel whether it is part of the cell or not, appeared to be more difficult than the recognition of RBCs. As intended, cells produced a strong signal, whereas most floating artefacts were ignored unless they came into direct contact with the cells (Figure 3b, eighth frame, bottom row). Such temporary associations, however, have minimal effects on the ΔDI calculations (Methods). The tracking script was tasked with ignoring cells that travelled in clusters or collided with other cells during the journey. This was to ensure that the collision effects would not interfere with later DI calculations. As discussed in the synthetic data section, we trained the CNN on round and bullet-like shaped cells, and, as expected, it was able to correctly identify and segment similar real-world cells (Figure 3a). Moreover, the CNN was able to generalise to cells with abnormal shapes and textures, which are common for malaria iRBCs, as can be appreciated from Figure 3c for which we selected some striking ‘edge’ cases of cells with abnormal shapes. Since the CNN output contained the position of cells as peaks of probability values, we could correct for false positives and false negatives by setting a threshold to the model’s output. We observed that a threshold for recognising a cell that was set too high led to missing cells that were almost transparent in some of the frames. A too low threshold, however, returned an increasing number of false-positives, typically large chunks of cell debris or parasite clusters (e.g., Figure 1d, bottom row). We preferred false positives over false negatives since we did not want to overlook any true cells, and a visual inspection of the cell journeys allowed us to remove the obvious non-cells afterwards. The inspection also revealed unexpected ‘double-cell’ journeys overlooked by the tracking algorithm caused by the CNN recognising two overlapping individual cells as one single cell (e.g., the top row of Figure 3d).

**Figure 3.**
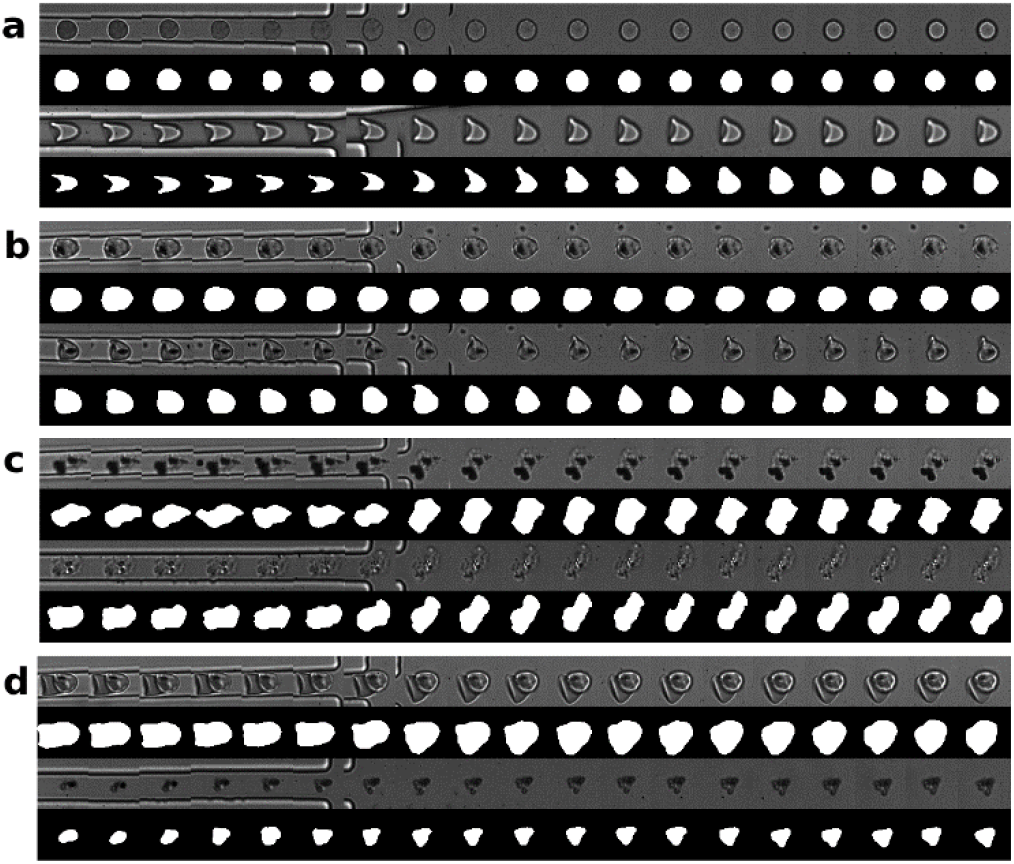
Cell tracking results. Original images are on top, segmentation results are at the bottom. (a) Examples of cell-journeys of cells with shapes similar to those in the training dataset. (b) Examples of cell-journeys with noise artefacts floating around the cell, which could potentially hinder correct cell-segmentations. Note that in the in the 8th frame of the bottom journey, the artefact is in contact with the cell and becomes part of the cell-segmentation. (c) Cell journeys with unusual shapes not similar to the training data. This demonstrates the generalization abilities of the neural network. (d) Some ‘failure’ cases. The top journey contains two cells traveling together, which should have been excluded. This was probably caused by the fact that they were overlapping the whole journey making it look like a single cell to the neural network. The bottom one is debris that was seen as a cell by the neural network due to its large size.

### Quantitative comparison of annotations

Although we visually confirmed the neural network performed adequately in segmenting the cells, we also wanted to assess the results quantitatively. To assess the CNN’s performance, we compared the RBC surface area as determined by three human curators with experience in analysing RBCs (GT, JK, and DR) with that of the CNN. Curators each were given samples from the 81,034 unique cell pictures segmented by the CNN (cropped to 54×54 pixels) without its output-prediction and were instructed to segment them by drawing polygons around them to indicate cell form. Agreements between shapes were quantified as fractions of overlapping pixels. As ambiguous or ‘edge cases’ are relatively hard to segment, we made sure they were well represented in the set (∼20%). To get an indication of the overlap between the human and CNN segmentations, we also compared them among curators themselves. We found that, collectively, the human curators had a median overlapping fraction of .950 with the CNN’s output, and a median fraction of .961 amongst themselves (Table 1). Given that the cells are centred in the images, we actually expected large portions of the labels to always overlap. As a sanity check, we also evaluated what these fractions would be if the CNN outputs were compared to randomly selected curated segmentations of the cells. When randomly paired, the median overlapping fraction drops to ∼.90 (Table 1, column 4). The small difference between human and CNN assessment, as well as the visual inspection of the CNN’s outputs, gave us confidence in the CNN’s ability to segment cells. Note that for the final ΔDI determination (Figure 4), we only used the median of the DI values between the shapes during and after the constriction (Methods). This was done to ensure that occasional mis-segmentations (e.g., caused by cells being too transparent or touching floating cell debris) did not play a large role in the ΔDI calculations. These potential missegmentations were however not excluded from the samples on which Table 1 is based, allowing for an unbiased assessment of the CNN’s capabilities. As a result, we anticipate that the final segmentations used for ΔDI calculations will even be more similar to the human curator results than Table 1 indicates.

**Figure 4.**
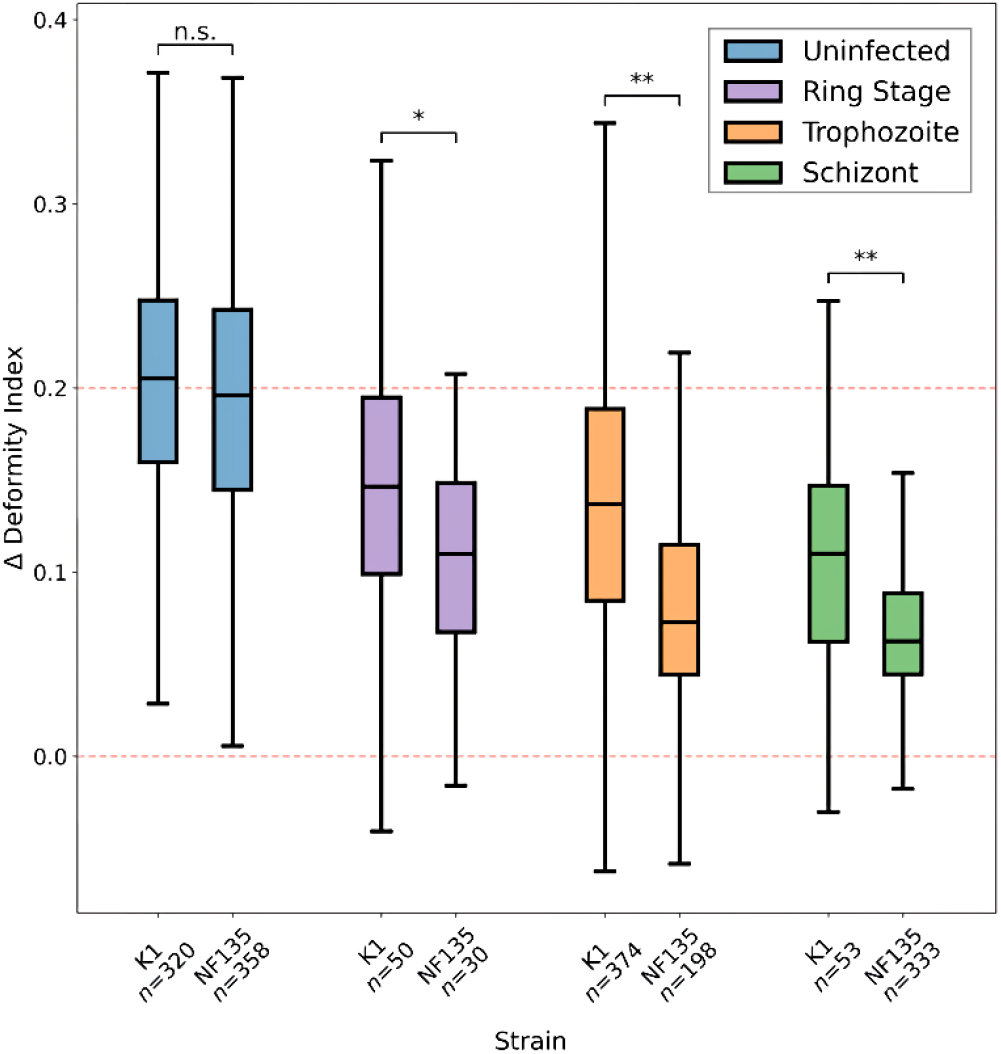
ΔDI distributions of the various infection stages, per strain. Student’s t-test between each infection stage of both strains. The significance of the Student’s t-test is indicated as follows: n.s. = P > 0.01;* = 0.01 > P > 0.001;** = P < 0.001. The boxplots contain a middle line that indicates the median; the lower and upper ends of the box indicate the 25th and 75th percentiles; and the lower and upper extremities indicate the minimum and maximum values. The dotted red lines indicate the zero-deformity and the median-deformity of uninfected cells.

**TABLE 1.** Trained CNN assessment

| Curators | Overlapping Samples | Surface area median fraction overlapping | Surface area median fraction overlapping, randomly paired |
| --- | --- | --- | --- |
| Curator1 - CNN | 105 | .947 ± .0384 | .895 ± .0437 |
| Curator2 - CNN | 500 | .953 ± .0464 | .893 ± .0421 |
| Curator3 - CNN | 500 | .947 ± .0457 | .901 ± .0562 |
| All curators - CNN | 1105 | .950 ± .0454 | .896 ± .0464 |
| Curator1 - Curator2 | 105 | .973 ± .0122 | .898 ± .0409 |
| Curator1 - Curator3 | 105 | .949 ± .0243 | .902 ± .0431 |
| Curator2 - Curator3 | 500 | .960 ± .0256 | .904 ± .0425 |
| All curator combinations combined | 1105 | .961 ± .0290 | .904 ± .0401 |
| CNN - CNN | 81034 | 1.00 ± .000 | .895 ± .0581 |

### The effect of developmental stage and strain on deformability

Having validated our synthetic data approach for training the CNN, we next focused on studying the effect of malaria strain and developmental stage on the ΔDI of iRBCs. In this study we compared two P. falciparum strains of different geographical origins: K1 from Thailand and NF135 from Cambodia. For both strains, we studied the parasite effect in the ring, trophozoite, and schizont developmental stages on the ΔDI and used uninfected RBCs as controls for maximally deformable cells. In total, 1,716 cells were evaluated.

Infected RBCs have a lower ΔDI than non-infected ones, and the ΔDI is lowest in the largest and most mature developmental stage. Interestingly, we also find a significant and consistent difference across the developmental stages between the two strains, with NF135-infected RBCs having a lower deformability than K1-infected RBCs, while there is no difference between the ΔDI of the uninfected RBCs.

## 3. Discussion

We described a deep learning approach that enabled us to investigate the effects of malaria strain and phase on the deformability of infected RBCs. To accomplish this, we used synthetic data to train a CNN to predict RBC surface areas and locations and process that information with a Python script. We demonstrated that the neural network performed well in noisy environments and could generalise not only to real-world data, but also to cells with abnormal shapes and textures. We found a significant RBC deformability difference between the K1 and NF135 strains throughout asexual blood-stage development. To the best of our knowledge, we are the first to use deep learning for automatic RBC deformity quantification under noisy conditions and the first to use non-simulation synthetic data to solve this problem.

The observed difference in the deformability between the strains is particularly interesting as differences in the membrane rigidity of iRBCs have been linked to virulence in the asexual blood stages of the parasite and to the transmission of the sexual blood-stage parasites to the mosquito vector[5]. The variation between strains in the asexual blood stages tested here could be indicative of natural variation in parasite virulence[2]–[4]. While disease severity has been reported for the K1 strain[24], [25], there is no data on the disease severity of NF135. Although controlled human malaria infections (CHMI) with the NF135 strain in naive individuals did not report severe malaria during the relatively short infection period[21, p. 135], direct patient comparisons would be required to confirm the hypothesis that the differences in the ΔDI observed between the strains correlate with differences in virulence. However, the increased membrane rigidity combined with the drug resistance profiles of NF135[21, p. 135] could make this particular parasite strain vital to uncovering important molecular mechanisms relevant to disease progression and parasite survival within the human host. Regardless of the interest surrounding NF135, the observed differences in membrane rigidity between these two wild-type malaria strains using this technique underline its power to uncover discrete differences in membrane mechanics. This will undoubtedly aid in the comparison of strains as well as genetically modified parasites in the quest to unravel the molecular mechanisms of virulence. Furthermore, by applying the same strategy to studying sexual bloodstage deformability, the importance of the relative levels of RBC rigidity for transmission can be explored. Finally, this technique can be used to study existing compounds and discover new ones aimed at reducing rheological changes in the iRBC that the current filtration-based screen would miss [26].

It is well established that more diverse training data leads to more accurate DL models because deep learning can only interpolate between what it has observed during training[27], [28]. As a consequence, DL often fails on edge cases, which are rare occurrences that are unlikely to be captured in a training dataset. With our synthetic data, we were able to add rare but critical edge situations purposefully at any desired quantity and with consistent labelling. We found that by exaggerating the data features, the neural network learned to generalise better to new data. Also, by training on synthetic data while testing on real-world data, we did not have to be concerned with data leakage between training and test sets, something that is becoming a major part of the DL replication crisis in many science-fields[29]. However, there are also valid arguments against the use of synthetic data to answer research questions. The most important one being that synthetic data will always be based on a rough approximation of the real world and thus also be limited by that. This could potentially result in false insights and, as a result, incorrect decision-making. After all, you can only get as much out of the data as you put in. What distinguishes this case is that the employed deep learning model is not attempting to learn something novel from the data but rather learns to take over the mundane task of cell segmentation under noisy conditions. The synthetic data used in this study is more than adequate for that task. The CNN is only used for what it is good at, namely pattern recognition, and statistical analysis is used to detect meaningful patterns in the data. This approach allows researchers to focus on their own strengths, such as hypothesis generation and testing, instead of wasting time labelling data by hand.

The cell detection and segmentation procedures took care of the majority of the labour involved in analysing MD videos. Yet, the extracted cells still had to be categorised into the malaria stages manually. A natural next step could be to generate synthetic data representing all malaria stages so that a neural network could also learn to classify those. However, given the significant variance introduced by the random generators during data generation, which makes it difficult to replicate the small distinctions between malaria stages, as well as the method’s simplicity, we do not feel this strategy is viable. Instead, we propose that the synthetic data be limited to cell tracking and extraction alone. We believe that manually browsing and classifying the successfully processed cell journeys with the corresponding segmentation results even has its advantages. For starters, it allows researchers to define their own thresholds between malaria stages. Also, it enables researchers to disqualify obvious mistakes made by the programme (e.g., Figure 3d), which benefits science by increasing trust in the results.

Besides this study’s objective, the technique discussed here is relevant to more than just studying malaria-induced RBC effects. In the future, this analysis pipeline could be used to study RBC deformability in diseases such as sickle cell anaemia, thalassemia, and hereditary spherocytosis, to name a few. This tool can also be used to study RBC dynamics during other physiological processes, such as hypoxia, oxidative stress, lipid peroxidation, and the effects of red blood cell membrane drug binding. And these are only a few RBC-specific examples. We expect that by modifying the synthetic data, it can be extended beyond analysing RBCs to other single cells or droplets, but this would require further research to confirm. Thus, the proposed pipeline has the potential to open up a whole new avenue of study.

To conclude, by showcasing the success of DL in one of the more difficult MD cases, we demonstrated that DL has the potential to significantly accelerate the analysis of single-cell studies that use microfluidic devices. We found that, in some cases, such as in this study, synthetic data can effectively complement DL. We anticipate that our approach will make the benefits of DL, which were previously restricted to manually labelling data, available to a larger audience. However, we also believe that DL should be employed conservatively to promote research transparency, which can only benefit science. The newly discovered effect of malaria strain on the deformability of iRBCs suggests that our combination of DL with CNNs and synthetic data can be used to uncover factors underlying the reduced deformability of iRBCs in malaria.

## 4. Methods

### Microfluidic devices and experimental protocol

A microfluidic device consists of an inlet, 30 parallel channels (6 regions with 5 channels in each), and an outlet (see Figure 5). A narrow channel was 7 μm wide and 1 mm long, two adjacent channels were 13 μm apart. A microfluidic pattern was drawn in Autocad (Autodesk) and transferred to glass (JD Photo Data). A microfluidic master was fabricated by patterning SU8 2007 photoresist (Kayaku Advanced Materials) on a silicon wafer (50 mm dia.; Si-mat); the photoresist was processed according to the manufacturer’s guidelines. Briefly, photoresist was spun (20 seconds at 500 rpm; 30 s at 3000 rpm; acceleration 300 rpm/s) on a wafer, baked at 65 °C for 1 minute, and at 95 °C for 2.5 minutes; To define the pattern, the wafer was exposed through a glass mask (dose 100 mJ/cm2; mask aligner MBJ3, Suss Microtec), baked at 65 °C for 1 minute, and at 95 °C for 3 minutes. The pattern was developed in SU-8 developer (Kayaku Advanced Materials) and the obtained master was baked at 175 °C for 2 minutes. The height of fabricated features was checked with a Dektak 6M stylus profiler (Bruker) and was between 8.5 and 9 μm. The surface of the master was treated with 1H,1H,2H,2H-perfluorooctyltrichlorosilane (Thermo Scientific) to promote the removal of elastomer; the master and 50 μL of silane were left in a desiccator for 1 h under vacuum, followed by 2 h in a 95 °C oven.

**Figure 5.**
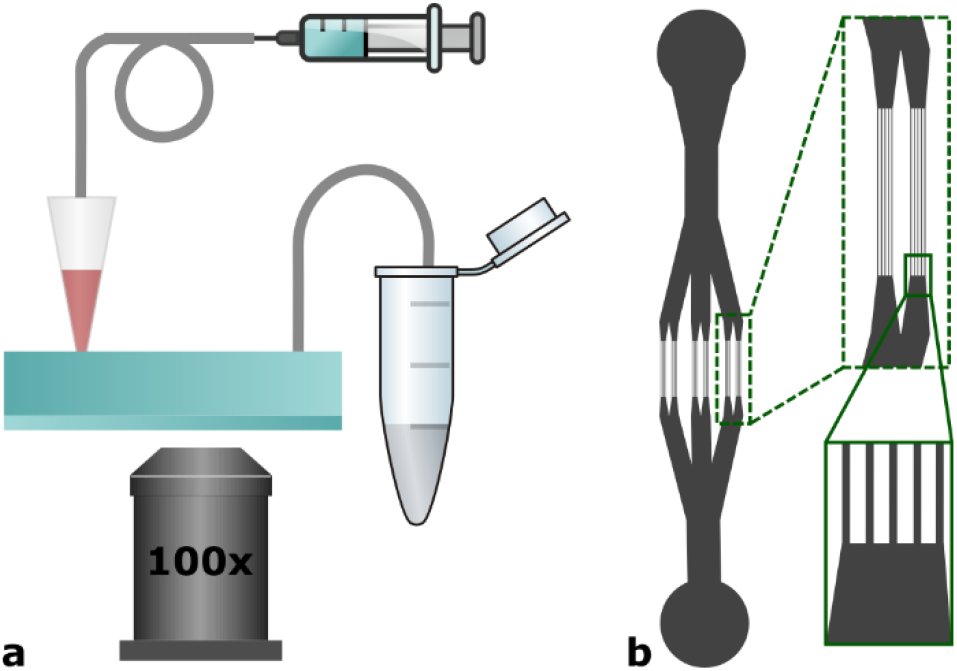
An experimental setup to capture RBCs deformation. (a) Overview of a setup: a syringe with sacrificial fluid (HFE7500) and a 200 μL pipet tip is connected to the inlet of a microfluidic device. A pipet tip is loaded with 50 μL of RBC sample. An outlet is connected to a waste vial. Deformation of RBCs is observed on the IX71 inverted microscope via a 100x oil immersion objective. (b) An architecture of a microfluidic device: the inlet (circle at the top) splits into 6 sets of cannels; each set consists of five 7 μm narrow and 1000 μm long channels. Neighbouring channels are 13 μm apart. The outlet is located at the bottom. The inset at the bottom-right of the panel shows the approximate position where experimental videos were acquired.

Microfluidic devices were made from Sylgard 184 Silicon Elastomer kit (Dow). Base and cross-linker parts were mixed in a ratio of 10:1 (w/w), poured over the master to create a 5-7 mm thick layer, and degassed in the desiccator. PDMS was cured for at least 2 h at 65 °C in an oven. PDMS was separated from a master, and a biopsy punch (1 mm dia., Kai Medical) was used to bore 1 mm holes for the inlet and outlet. PDMS piece was cleaned with Scotch tape, rinsed with isopropanol, and blow dried with nitrogen gas. Finally, a glass coverslip (50 mm dia.) and the PDMS piece were treated with oxygen plasma (25 s, 65 W, Femto 1A, Diener Electronic), and sealed together.

An experimental setup consisted of an IX71 (Olympus) microscope equipped with a 100x oil immersion objective (UPlanFLN, Olympus) and Miro ex4 (Vision Research) camera. Sample flow was controlled by Nemesys low-pressure syringe pump (Cetoni).

Ethanol was used to fill the narrow and shallow channels of a device and then replaced with PBS before introducing a sample.

A PDMS connector: a 5 mm biopsy punch (Kai Medical) was used to cut out a PDMS cylinder from a 5-7 mm thick PDMS sheet. 1 mm hole was bored in the centre of a PDMS cylinder along the rotational symmetry axis.

A 0.5 mL Gastight syringe (Hamilton) was filled with HFE7500 oil and connected to tubing (0.56 mm ID, 1.07 mm OD, Adtech Polymer Engineering) using a 23G (blue) needle. A PDMS connector was used to connect tubing to a 200 μL pipet tip. Prior to loading the sample, a pipet tip was filled with HFE7500 oil from a syringe. 50 μL of a sample was loaded into a tip and connected to a microfluidic chip; the other outlet was connected to a piece of tubing and a waste vial.

A sample aliquot was pushed through a device at 50 to 100 μL/h. Movies were acquired at a rate of 1000 fps.

### Malaria data

Plasmodium falciparum strains K1 and NF135 were cultured in standard culture conditions[30] in RPMI media supplemented with 10% human serum and maintained in 5% haematocrit of human RBCs. Samples for microfluidic analysis were taken directly from the cultures and kept at 37 °C until they were directly added to the microfluidic device.

### Synthetic RBCs

As illustrated in Figure 6, the synthetic data consisted of images of 100×100 pixels with a single-colour (grayscale) dimension. Each image consisted of a noisy background (e.g., Figure 6a, panel 2) with some or all of the following objects: a single wall (Figure 6a, panel 3), static artefacts (Figure 6a, panels 4 and 6), floating objects (Figure 6a, panel 5), and a round (Figure 6b) or bullet-like shaped cell (Figure 6c). Besides drawing circles and lines, the main algorithms used are: elastic deformation[22] (Figure 6d), Gaussian distribution function (examples in Figure 6e), and simplex noise generation[23] (e.g., Figure 6f). Elastic deformation is a function that can ‘randomly’ warp images, the ranges of which are determined by the parameters α and σ, which determine the range and intensity of image-warping. The Gaussian function creates a two-dimensional bell-curve. It contains a single parameter, the standard deviation (σ). The σ can either be used directly or be calculated indirectly given a box-size (e.g., Figure 6e had box-sizes of 3 and 16 and an ideal σ was calculated for those sizes). The OpenSimplex function, which generates gradient-noise patterns, has three parameters: the zoom, which determines the ‘size’ of the noise (for example, the left image in Figure 6f has a smaller zoom value than the right image), the random seed, which serves as a starting point for the random number generator used by the algorithm, and the range, which normalises the noise to be within a certain range of values. The seeds were based on the local time. Because we employ a large number of randomly sampled integers to construct the synthetic data, these will be designated as rdm_range(n,m) in the remainder of the article, signifying any integer randomly chosen between n and (including) m. Often, multiple images were interpolated (combined) into a new image. Interpolation requires two images (P1 and P2) and a third interpolation image (I) that contains information on how much each pixel in either image should contribute to the new combined image (P3) and has only values ranging between 0 and 1. Interpolation is then done using the formula:

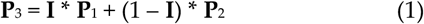

**Figure 6.**
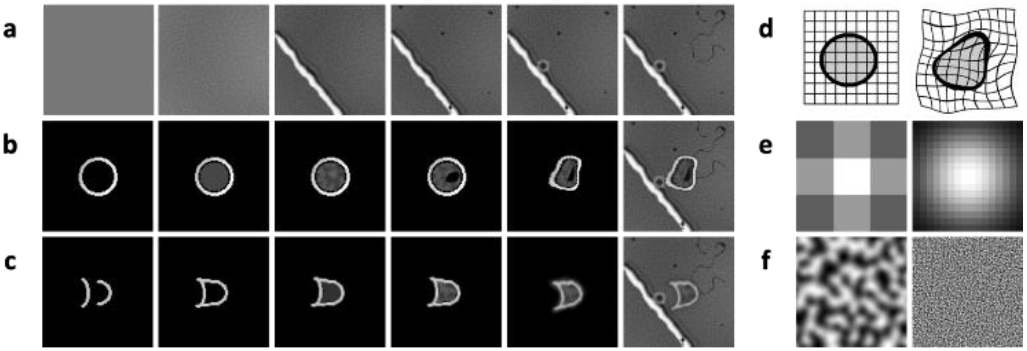
Creation of synthetic RBCs. (a) Steps in the creation of synthetic backgrounds (b) Steps in the creation of synthetic round cells. c Similar as a, but for bullet-like cells. (d) Illustration of elastic warping, which was used to introduce variation to object shapes. (e) Gaussian distribution examples, which were used to smooth out rough edges (left), but also to generate the cell’s centre-probability labels (right) (f) Some examples from the OpenSimplex noise function. These are gradient noise patterns that can be utilised to synthesise textures (as seen in a’s second panel, b’s third panel, or c’s fourth panel), but they can also be used to interpolate between two images, with the intensity of the noise value determining how much each of the two images per pixel should contribute to the combined image (e.g., a noise value for a given pixel of .7 corresponds to the combined pixel being 70% image 1 and 30% image 2 for that pixel).

A base background (Figure 6a last panel) always started from a colour=rdm_range(65, 146) and added Simplex noise with zoom=1 and range=(0, 1). The image was then blurred with a Gaussian kernel with box-size=3×3. After-wards, another Simplex noise-pattern with zoom=1 and range=(–15, 15), was added. These steps created the small fine noise (e.g., Figure 6a 2nd panel). Then, another Simplex noise-pattern with zoom=0.01 and range=(−10,10) was also added to give the images some colour gradient (e.g., Figure 6a 3rd panel). The addition of a wall object was only done for 60% of the images. Wall objects were made by drawing 3 lines with colours rdm_range(85, 140), rdm_range(0, 15) and rdm_range(200, 255) and a thickness of rdm_range(1, 4) pixels for the first two lines and rdm_range(3 and 7) pixels for the last line. The drawn lines were then rotated randomly (together). About 40% of the images with a wall object were warped with elastic deformation with α=700 and σ=14. The remainder has an elastic deformation with α=15 and σ=3. Finally, the wall object was blurred with a Gaussian kernel of 5×5 pixels and placed in the base background image (e.g., Figure 6a 3rd panel). Artefacts, as can be seen in Figure 6a’s 4th panel, were created seven-fold. They were made by drawing small black (colour=0) filled circles with a radius of range=(1, 3) pixels, and warping them with elastic-deformation with α=300 and σ=12 before placing them in the base background on random locations. Floating objects (e.g., Figure 6a 5th panel), were created in 66% of the images. They were made by drawing two circles: one with a radius of rdm_range(2,5) pixels and the other with rdm_range(0,3) pixels. The colours for both circles were chosen to be either from rdm_range(200, 255) or from rdm_range(28,38). After drawing, they were blurred with a Gaussian kernel of 5×5 pixels and placed in the base-background image at a random position. The black, hair-like artefact, as can be seen in the 6th panel of Figure 6a, was created by drawing a black line with a length of rdm_range(0, 25) pixels, and a thickness of rdm_range(1, 3) pixels. This artefact was than warped with elastic deformation with α=500 and σ=6 and placed in the base-image at a random position and random rotation. A second base background, with all objects besides floating objects or cells, was also generated with a 50% chance of being lighter or darker. This to simulate the median-frame the neural network also receives as input.

Round blood cells started with a drawn circle with radius=16 pixels, thickness=rdm_range(1,4) pixels, and colour=rdm_range(110, 250). Then, a second circle was drawn with radius=14 pixels, thickness=rdm_range(1,3) pixels, and colour=0. After that, a filled circle was drawn with radius=13 pixels and colour=70 (e.g., 2nd panel of Figure 3b). The texture of the inner cell (e.g., the 3rd panel of Figure 6b) was created by replacing the inner circle with a Simplex noise pattern with zoom=0.2 and range either (50,100) or (0,220). A malaria parasite was simulated by drawing a warped black dot in the cell, warping was done with elastic deformation with α=300 and σ=12 (e.g., the 4th panel of Figure 3b). After adding the parasite, the whole cell was warped with elastic deformation with α=200 and σ=either 8 or 12 (e.g., 5th panel of Figure 6b). The cell was then blurred with a Gaussian kernel of 3×3 pixels, and added to the centre of the base background (e.g., Figure 6b’s 6th panel) via interpolation. Bullet cells started a bit differently. They started with two half circles with different curvatures and thickness=2 (e.g., Figure 6c 1st panel). The centres were rdm_range(−20,10) pixels apart. The edges of the circles were connected with lines that were also extended on the left side with rdm_range(0,5) pixels (to simulate tail extension, e.g., Figure 6c’s 2nd panel). The cells were then either filled uniformly with colour=rdm_range(85, 140) (e.g., 3rd panel of Figure 6c) or filled with a texture (e.g., 4th panel of Figure 6c). This texture was made of Simplex noise with zoom=.2 and range of either (50, 100) or (0, 220). The bullet cell was then blurred with a Gaussian kernel of size (9,9), warped with elastic deformation with α=400 and σ=17, rotated with an angle of rdm_range(−30, 30) degrees (e.g., Figure 6c, 5th panel), and then placed in the base background via interpolation (e.g., Figure 6c, 6th panel).

### Expert labelling of real RBCs

A simple tool was created to aid in the manual labelling of retrieved RBCs. A domain expert was given 2.461 cell journeys to label via this tool. This, with the following categories: 1) uninfected; 2) ring stage; 3) trophozoite; 4) schizont; and 5) other. All images were shown without context (e.g., the source of the RBC cell journey) and shuffled at random prior to the classification process.

### Convolutional neural network architecture

Inspired by the human ventral and dorsal visual pathways, we designed the neural network to include two output layers: one for segmentation to determine cell shapes, and one for detecting RBCs and their locations. There are seven hidden layers in the CNN. The first layer has a dilation parameter of 30, which adds a black border of 30 pixels around the input. Figure 7 shows that the dilation of the hidden layers goes up and down[31], which makes the receptive field bigger without the need for pooling layers. The kernel size is 5×5 for all layers, including the output layers. Batch normalisation and ReLU activation were applied to all hidden layers. The output layers have a Softmax activation function for the cell-segmentation output and a Sigmoid activation function for the cell-location output. Since no tensor flattening was used, this neural architecture design is able to handle larger inputs than it was trained on.

**Figure 7.**
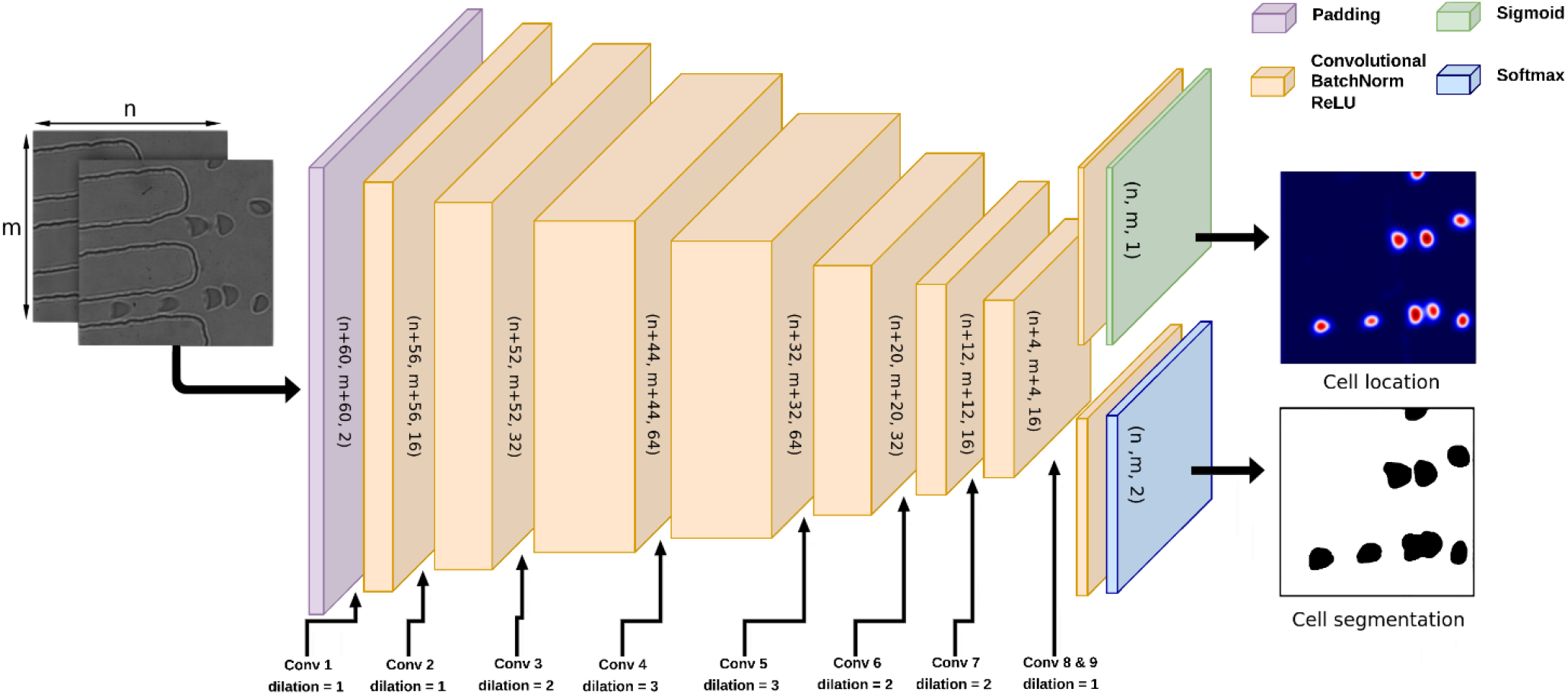
Neural network architecture for segmentation and instancing of RBCs. The neural network consists of nine convolutional layers. The first layer adds padding around the input image, and the layers three through seven have an increase and then a decrease in dilation. The last two layers only consist of a convolutional layer and an activation function. Sigmoid was used for the instancing and Softmax for the segmentation.

### Training, validation, and testing procedure

A model based on the aforementioned architecture was trained on 10,000 synthetic data instances with an equal number of round and bullet-shaped cells. We trained on minibatches containing 32 samples each. The binary cross-entropy loss was used to calculate the loss between the CNN outputs and the labels. The loss values for both types of labels were summed. The Adam optimizer[32], with the standard learning rate of 10-3, was used to optimise the model parameters. The model parameters were saved with every ten minibatches of training. Training was automatically stopped after 22 epochs.

The validation set was a single frame from 2 videos used in this study (K1_001_20201105.tif, and NF135_002_20201105.tif), and an unrelated mouse-RBC MD videoframe (002_2.5kfps.avi). The mouse video frame was selected to evaluate the model’s generalisation abilities. The three frames contained 38 cell images in total. The optimal parameters were chosen by visually examining the model’s output from the three validation frames for each of the saved checkpoints. We refer to the GitHub page for the actual validation frames and the Zenodo webserver for all the videos.

We used data from all videos to test the segmentation output quality. First, the model was applied to all movies, yielding 2,461 cell-journeys with approximately 33 sub-images each. Then, these cell sub-images were cropped from video frames with a size of 128×128 (with the cell exactly in the middle), yielding 81.034 images of single cells. A Python programme was created to assist with manual labelling. Human experts were randomly given single images of cells by the programme. These images were four times enlarged, and presented to annotators without any knowledge of the model’s output. The programme enabled them to draw a polygon around the cell to define its form. The outputs of both human curators and the CNN were cropped to 54×54 pixels and compared (Table 1).

### Cell tracking script

The cell-tracking programme, written in Python, was designed to track individual cells through the MD and to calculate the ΔDI. The script used the CNN’s output as its input. The tracking itself only requires the cell-location output of the neural network. Since collisions between cells could negatively influence the deformity determination, colliding cells and cells that travelled in clusters were excluded from ΔDI analysis by the programme.

Prior to processing, a ‘median frame’ was created for each MD video by selecting 100 frames at random from a specific movie and calculating the median value for each pixel. The videos were split and processed in parallel in order to improve analysis time. Each movie was divided into 20 chunks, with 60 frames overlapping between consecutive chunks. Each frame was concatenated with the median frame to meet the CNN’s input dimension requirement. Feeding the concatenated frames to the model resulted in the segmentation (representing RBC shapes) and location-peak predictions for each frame. Per frame, all location-peak outputs were smoothed with a Gaussian-blur function with a kernelsize of 3×3. The peak-centres with probability values larger than 0.3 and at least 10 pixels apart from other peaks were considered to potentially represent the centre (x,y) coordinates of cells. A ‘cell initialization threshold’ was set at 70px, illustrated as a line in Figure 8. All cells detected left of this threshold were considered as ‘new entering cell objects’. For all new cell objects, frame crops of 64×64 pixels around the middle of the cell objects, as well as a same-sized crop around the CNN segmentation output, were stored in memory. To enable cell tracking, each cell object’s next cell location was first predicted before processing the next frame. This was accomplished by adding the velocity of the cell to the position using the formula:

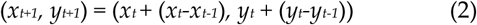

**Figure 8.**
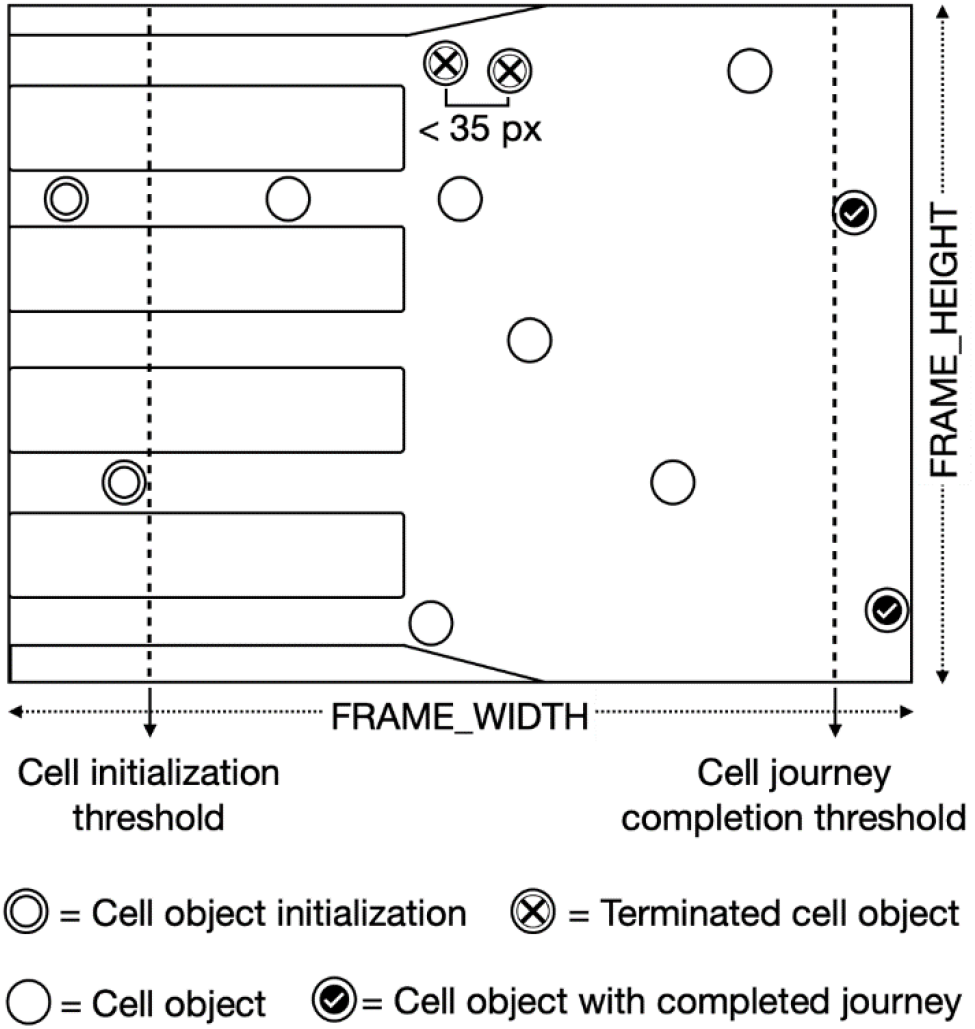
A schematic of the tracking algorithm. A cell initialization threshold indicates the start of a cell-journey. With every consecutive frame, the cell’s new coordinates are predicted given the cell’s momentum. Cells are eliminated for research if they come within 35 pixels of another cell. After a cell object passes the cell journey completion threshold it is marked as complete

These predictions were then compared to the CNN’s cell-location coordinates of the next frame to check if they were similar. A match was defined as a Euclidean distance that was less than twice the cell velocity and no larger than 30 pixels. Cell-objects that did not have a match with the location peaks were removed from memory. The crops of the frame and segmentation results were stored in memory alongside earlier crops for the cells that did match. The distances between all cell-objects were then calculated, and cells with a distance of less than 35 pixels were erased from memory to ensure colliding cells were not part of the study. The process of predicting the next location of cell objects and matching them with CNN detections was repeated until the cell-objects met the ‘completion threshold’, which was defined as the frame-width minus 40 pixels (see Figure 8). Following that, all cell-crops were stored as cell-journey images (e.g., Figure 2).

### Shape and Deformity Index determination

The deformity of an RBC is defined as the difference between the RBC’s width (W) and height (H) ratios during and after constriction (formula 3).

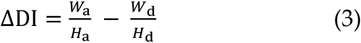

Each RBC, however, has many frames during the constriction and multiple frames after the constriction. Therefore, we calculated the DI for each frame and used the median DI value of the constriction and the median DI value outside the constriction to calculate the ΔDI. The rationale behind this was that this would result in a more robust ΔDI since noise factors, like those illustrated in Figure 3B, could sometimes hinder making proper segmentations.

Before calculating the DIs, cell-journeys with raw segmentation CNN outputs were multiplied by 255 and saved as grayscale images (grayscale = 1 colour dimension per pixel instead of the 3 used in colour images). For DI calculation, the cell journey segmentation outputs were loaded in and first smoothed with a Gaussian function (σ value of 1) and then binarized using a threshold of 150. Some frames of the journeys still contained parts of nearby cells. Therefore, a ‘Flood Fill’[33] function was applied to all frame centres to isolate the cell segmentations from other cells. Afterwards, rectangles were drawn around all cell segmentations to create bounding boxes. The widths and heights of the boxes were used to compute the cell ratios.

## Data and code availability

The microfluidic device videos have been deposited on the Zenodo webserver.

(https://doi.org/10.5281/zenodo.7374342)

The code used to generate the synthetic images, train the CNN and process all results are available at: https://github.com/cmbi/synthetic_cells

## Acknowledgsments

D.R. acknowledges the financial support provided by the Hypatia Fellowship from Radboudumc (Rv819.52706). N.P. was supported by a Marie-Sklodowska Curie grant (790085) and T.K. and G.T. by the Netherlands Organisation for Scientific Research (NWO-VIDI 864.13.009)

Special thanks to Alex van der Starre, Julie Verhoef, Cas Boshoven, and Felix Evers for their assistance in evaluating the domain experts’ image classification labels.

The authors thank Joshua Gillard for assisting in the graph design of Figure 4. We would also like to thank dr. Li Xue, the direct supervisor of D.R., for facilitating this project.

## Author contributions

N.P. and T.K. conceived the strain-effect hypothesis on red blood cell deformability, designed the wet-lab part of the study, and provided the biological samples. D.R. conceived the deep learning and synthetic data approaches, wrote the first draft of the paper, created Figures 1,2,3 and 7, and designed the original code. J.K. revised the code and designed Figures 4, 6 and 8. A.P. and W.H. were responsible for handling the microfluidic devices and recording the data. A.P. also designed Figure 5. D.R., G.T. and J.K. participated in manually annotating cell shapes for Table 1. D.R. created the associated tool to do so. N.P. classified all the malaria stages. D.R., M.H., A.P., and G.V. contributed in writing the manuscript. All authors critically reviewed the manuscript.

## References

[1] World Health Organization, World malaria report 2021. Geneva: World Health Organization, 2021. Accessed: Aug. 09, 2022. [Online]. Available: https://apps.who.int/iris/handle/10665/350147

[2] T. K. Jonsdottir, M. Gabriela, B. S. Crabb, T. F de Koning-Ward, and P. R. Gilson, ‘Defining the Essential Exportome of the Malaria Parasite’, Trends Parasitol., vol. 37, no. 7, pp. 664–675, Jul. 2021, doi: 10.1016/j.pt.2021.04.009.

[3] B. M. Cooke, J. Stuart, and G. B. Nash, ‘The cellular and molecular rheology of malaria’, Biorheology, vol. 51, no. 2–3, pp. 99–119, 2014, doi: 10.3233/BIR-140654.

[4] N. I. Proellocks, R. L. Coppel, N. Mohandas, and B. M. Cooke, ‘Malaria Parasite Proteins and Their Role in Alteration of the Structure and Function of Red Blood Cells’, Adv. Parasitol., vol. 91, pp. 1–86, 2016, doi: 10.1016/bs.apar.2015.09.002.

[5] M. Tibúrcio et al., ‘A switch in infected erythrocyte deformability at the maturation and blood circulation of Plasmodium falciparum transmission stages’, Blood, vol. 119, o. 24, pp. e172–180, Jun. 2012, doi: 10.1182/blood-2012-03-414557.

[6] M. Aingaran et al., ‘Host cell deformability is linked to transmission in the human malaria parasite Plasmodium falciparum’, Cell. Microbiol., vol. 14, o. 7, pp. 983–993, Jul. 2012, doi: 10.1111/j.1462-5822.2012.01786.x.

[7] D. Bento et al., ‘Deformation of Red Blood Cells, Air Bubbles, and Droplets in Microfluidic Devices: Flow Visualizations and Measurements’, Micromachines, vol. 9, o. 4, p. E151, Mar. 2018, doi: 10.3390/mi9040151.

[8] H. Yin and D. Marshall, ‘Microfluidics for single cell analysis’, Curr. Opin. Biotechnol., vol. 23, no. 1, Art. no. 1, Feb. 2012, doi: 10.1016/j.cop-bio.2011.11.002.

[9] K. Matthews, E. S. Lamoureux, M.-E. Myrand-Lapierre, S. P. Duffy, and H. Ma, ‘Technologies for measuring red blood cell deformability’, Lab. Chip, vol. 22, o. 7, pp. 1254–1274, 2022, doi: 10.1039/D1LC01058A.

[10] J. C. A. Cluitmans, V. Chokkalingam, A. M. Janssen, R. Brock, W. T. S. Huck, and G. J. C. G. M. Bosman, ‘Alterations in red blood cell deformability during storage: a microfluidic approach’, Bio-Med Res. Int., vol. 2014, p. 764268, 2014, doi: 10.1155/2014/764268.

[11] G. Litjens et al., ‘A survey on deep learning in medical image analysis’, Med. Image Anal., vol. 42, pp. 60–88, Dec. 2017, doi: 10.1016/j.media.2017.07.005.

[12] A. Saadat et al., ‘A system for the high-throughput measurement of the shear modulus distribution of human red blood cells’, Lab. Chip, vol. 20, o. 16, pp. 2927–2936, 2020, doi: 10.1039/D0LC00283F.

[13] Y. LeCun, Y. Bengio, and G. Hinton, ‘Deep learning’, Nature, vol. 521, o. 7553, Art. no. 7553, May 2015, doi: 10.1038/nature14539.

[14] Z. Liu, H. Mao, C.-Y. Wu, C. Feichtenhofer, T. Darrell, and S. Xie, ‘A ConvNet for the 2020s’, 2022, doi: 10.48550/ARXIV.2201.03545.

[15] K. Simonyan and A. Zisserman, ‘Two-Stream Convolutional Networks for Action Recognition in Videos’, in Advances in Neural Information Processing Systems, 2014, vol. 27. [Online]. Available: https://proceedings.neurips.cc/pa-per/2014/file/00ec53c4682d36f5c4359f4ae7bd7ba1-Paper.pdf

[16] K. He, X. Zhang, S. Ren, and J. Sun, ‘Deep Residual Learning for Image Recognition’, 2015, doi: 10.48550/ARXIV.1512.03385.

[17] L. A. Gatys, A. S. Ecker, and M. Bethge, ‘Image Style Transfer Using Convolutional Neural Networks’, presented at the Proceedings of the IEEE Conference on Computer Vision and Pattern Recognition, 2016, pp. 2414–2423. Accessed: Aug. 09, 2022. [Online]. Available: https://openac-cess.thecvf.com/con-tent_cvpr_2016/html/Gatys_Image_Style_Trans-fer_CVPR_2016_paper.html

[18] Vijayalakshmi A and Rajesh Kanna B, ‘Deep learning approach to detect malaria from microscopic images’, Multimed. Tools Appl., vol. 79, no. 21–22, pp. 15297–15317, Jun. 2020, doi: 10.1007/s11042-019-7162-y.

[19] S. I. Nikolenko, ‘Synthetic Data for Deep Learning’, 2019, doi: 10.48550/ARXIV.1909.11512.

[20] N. N. Rahman, ‘Evaluation of the sensitivity in vitro of Plasmodium falciparum and in vivo of Plasmodium chabaudi Malaria to various drugs and their combinations’, Med. J. Malaysia, vol. 52, o. 4, pp. 390–398, Dec. 1997.

[21] A. C. Teirlinck et al., ‘NF135.C10: A New Plasmodium falciparum Clone for Controlled Human Malaria Infections’, J. Infect. Dis., vol. 207, o. 4, pp. 656–660, Feb. 2013, doi: 10.1093/infdis/jis725.

[22] P. Chlap, H. Min, N. Vandenberg, J. Dowling, L. Holloway, and A. Haworth, ‘A review of medical image data augmentation techniques for deep learning applications’, J. Med. Imaging Radiat. Oncol., vol. 65, o. 5, pp. 545–563, Aug. 2021, doi: 10.1111/1754-9485.13261.

[23] D. S. Ebert, Ed., Texturing & modeling: a procedural approach, 3rd ed. Amsterdam ; Boston: Academic Press, 2003.

[24] P. Sondo et al., ‘Genetically diverse Plasmodium falciparum infections, within-host competition and symptomatic malaria in humans’, Sci. Rep., vol. 9, o. 1, p. 127, Dec. 2019, doi: 10.1038/s41598-018-36493-y.

[25] F. Ariey et al., ‘Association of Severe Malaria with a Specific Plasmodium falciparum Genotype in French Guiana’, J. Infect. Dis., vol. 184, o. 2, pp. 237–241, Jul. 2001, doi: 10.1086/322012.

[26] J. Duez et al., ‘High-throughput microsphiltration to assess red blood cell deformability and screen for malaria transmission–blocking drugs’, Nat. Protoc., vol. 13, o. 6, pp. 1362–1376, Jun. 2018, doi: 10.1038/nprot.2018.035.

[27] F. Chollet, Deep learning with Python, Second edition. Shelter Island: Manning Publications, 2021.

[28] P. Domingos, ‘Every Model Learned by Gradient Descent Is Approximately a Kernel Machine’, 2020, doi: 10.48550/ARXIV.2012.00152.

[29] S. Kapoor and A. Narayanan, ‘Leakage and the Reproducibility Crisis in ML-based Science’, 2022, doi: 10.48550/ARXIV.2207.07048.

[30] M. van de Vegte-Bolmer et al., ‘A portfolio of geographically distinct laboratory-adapted Plasmodium falciparum clones with consistent infection rates in Anopheles mosquitoes’, Malar. J., vol. 20, o. 1, p. 381, Dec. 2021, doi: 10.1186/s12936-021-03912-x.

[31] R. Hamaguchi, A. Fujita, K. Nemoto, T. Imaizumi, and S. Hikosaka, ‘Effective Use of Dilated Convolutions for Segmenting Small Object Instances in Remote Sensing Imagery’. arXiv, Sep. 01, 2017. Accessed: Nov. 28, 2022. [Online]. Available: http://arxiv.org/abs/1709.00179

[32] D. P. Kingma and J. Ba, ‘Adam: A Method for Stochastic Optimization’, arXiv, arXiv:1412.6980, Jan. 2017. Accessed: Jul. 28, 2022. [Online]. Available: http://arxiv.org/abs/1412.6980

[33] G. Law, ‘Quantitative Comparison of Flood Fill and Modified Flood Fill Algorithms’, Int. J. Comput. Theory Eng., pp. 503–508, 2013, doi: 10.7763/IJCTE.2013.V5.738.

